# Insights into the cooperative nature of ATP hydrolysis in actin filaments

**DOI:** 10.1101/319558

**Authors:** Harshwardhan H. Katkar, Aram Davtyan, Aleksander E. P. Durumeric, Glen M. Hocky, Anthony C. Schramm, Enrique M. De La Cruz, Gregory A. Voth

## Abstract

Actin filaments continually assemble and disassemble within a cell. Assembled filaments “age” as a bound nucleotide ATP within each actin subunit quickly hydrolyzes, followed by a slower release of the phosphate P_i_, leaving behind a bound ADP. This subtle change in nucleotide state of actin subunits affects filament rigidity as well as its interactions with binding partners. We present here a systematic multiscale ultra-coarse-graining (UCG) approach that provides a computationally efficient way to simulate a long actin filament undergoing ATP hydrolysis and phosphate release reactions, while systematically taking into account available atomistic details. The slower conformational changes and their dependence on the chemical reactions are simulated with the UCG model by assigning internal states to the coarse-grained sites. Each state is represented by a unique potential surface of a local heterogeneous elastic network. Internal states undergo stochastic transitions that are coupled to conformations of the underlying molecular system. The UCG model reproduces mechanical properties of the filament and allows us to study whether fluctuations in actin subunits produce cooperative aging in the filament. Our model predicts that nucleotide state of neighboring subunit significantly modulates the reaction kinetics, implying cooperativity in ATP hydrolysis and P_i_ release. We further systematically coarse-grain the system into a Markov state model that incorporates assembly and disassembly, facilitating a direct comparison with previously published models. We find that cooperativity in ATP hydrolysis and P_i_ release significantly affects the filament growth dynamics only near the critical G-actin monomer concentration, while both cooperative and random mechanisms show similar growth dynamics far from the critical concentration. In contrast, filament composition in terms of the bound nucleotide distribution varies significantly at all monomer concentrations studied. These results provide new insights into the cooperative nature of ATP hydrolysis and P_i_ release and the implications it has for actin filament properties, providing novel predictions for future experimental studies.

## INTRODUCTION

Actin is a major component of the eukaryotic cytoskeleton and has important functions in cell motility and division. The monomeric form, globular actin (G-actin), polymerizes into filamentous actin (F-actin), and associates with filament binding proteins to form dynamic filaments of various architectures that are meticulously regulated to perform these functions. Actin monomers are large single-domain proteins made of 375 amino acids, and they contain a bound nucleotide at their center. The non-polymerized G-actin in the cell is predominantly found in an ATP-bound state (1). Polymerization of actin is followed by actin-catalyzed hydrolysis of the bound nucleotide ATP, and the hydrolysis in F-actin subunits has been estimated to be > 40,000 times faster than in G-actin, owing to the structural changes that G-actin undergoes as it transforms into an F-actin upon polymerization, during which it becomes more planar (2–6).

An actin monomer is not symmetric, and the actin filaments are polar in nature. New actin monomers predominantly add at the “barbed” end of the filament, and have a faster rate of depolymerization at the “pointed” end (7). Depending upon the G-actin concentration in the local environment, a filament can either grow from both ends (high concentration), shrink at both ends (low concentration), or grow at barbed end while shrinking at the pointed end. At a particular “critical” concentration, these two rates are balanced, and filaments undergo treadmilling, whereby a constant filament length is maintained on average. Experimental evidence suggests that incorporation of a new actin monomer into the filament does not immediately induce ATP hydrolysis, and nor does hydrolysis have to occur for subsequent monomers to be added (8). It is also believed that the rate of exchange of nucleotides in a filament with those in solution is negligible (9, 10). Instead, ATP is predominantly incorporated into the filament through polymerization of ATP bound G-actin. The interplay of polymerization and hydrolysis results in a time-lag associated with the hydrolysis in the filament with respect to polymerization and hence results in an ATP-bound cap in filaments at high G-actin concentration (11, 12). In F-actin, the hydrolysis of ATP to form ADP is not direct, as was initially hypothesized (11), but rather proceeds as a fast conversion to ADP with a protein bound inorganic phosphate (ADP-P_i_), followed by a slow release of the inorganic phosphate to the solution (13–16). ATP hydrolysis occurs on a time-scale of seconds, while P_i_ release takes place over minutes (2, 7, 13, 14, 17).

There are multiple cation binding sites in actin with varying affinities that modulate the mechanical properties of the filament (12, 18–21). Additionally, the state of the bound nucleotide also strongly influences its mechanical properties. The persistence length of actin filaments decreases as the bound nucleotide changes from ATP to ADP, as shown from experimental measurements (22).

Bottom-up coarse-grained (CG) models of actin filaments that are constructed using a heteroelastic network (23) based on reference all-atom simulations have been found adequate in capturing actin filament mechanical properties as a function of the state of the nucleotides, even at a highly coarse resolution of four CG sites per actin subunit (24, 25). A CG model with twelve CG sites per actin subunit has been shown to capture several other important structural aspects of actin (26). Such models that use a particle representation with an associated pairwise effective interaction potential are, however, often limited in their ability to represent certain molecular changes, including chemical or structural changes, that cannot be represented at the resolution of the CG sites, even if such changes ultimately affect the system behavior at the resolution of the CG model. Moreover, the underlying atomistic simulations further limit the configurations that such CG models can explore, as only configurations sampled by the atomistic model inform the effective potential, “locking” the CG model to, e.g., a given nucleotide composition. An example of such configurational changes is the state of the nucleotide bound to actin subunits in a filament. Although the CG particle representation has been shown to capture many essential conformational changes in actin subunits conditional on the state of bound nucleotide, these CG models do not offer insight about the hydrolysis dynamics involved, primarily because neither ATP hydrolysis nor P_i_ release are realized in all-atom simulations, due to the time-scales involved.

In contrast to the CG particle representation, a bottom-up model can be constructed using a discrete state representation, characterized by instantaneous transitions between different configurational states; these models are termed Markov State Models (MSM) (27–30). Generally, the rates governing the behavior of MSMs are obtained through statistics derived from fine-grained allatom simulations. Bottom-up MSMs are traditionally only able to study systems for which statistics have been directly obtained (29, 31, 32): including other states or rates in the model requires the computational scientist to use additional knowledge to determine the modified rate coefficients.

In this work, we first utilized the emerging concept of the ultra-coarse-grained (UCG) model, which combines both the particle and discrete state representations in a systematic way by defining an internal state associated with the CG particles (33–36). The internal states of the CG particles can in principle account for any reactions or conformational changes within the CG particles, making the UCG model ideal for actin filaments to study ATP hydrolysis and P_i_ release reactions. The UCG model was then used to systematically parametrize an MSM, which was analyzed to make conclusions about the spatial cooperativity present in an actin filament.

The macroscopic rates of the polymerization, de-polymerization, ATP hydrolysis and P_i_ release reactions have been measured indirectly from experiments using fluorescence labelling, radioactive labelling, etc., by assuming an underlying kinetic model (9–11, 13, 17, 37, 38). In experiments involving a conserved system, where the total mass of actin (G-actin + F-actin) in the system remains constant throughout the experiment, a characteristic sigmoidal shaped curve is obtained for time evolution of filament growth and ATP hydrolysis, showing that hydrolysis lags behind the polymerization at high initial G-actin concentrations (13, 39). Since it was established that ATP hydrolysis is mechanically decoupled from polymerization, there have been two major classes of hydrolysis models in the literature: the random model and the cooperative model. The random hydrolysis model (11, 31), first proposed before the intermediate ADP-P_i_ was discovered, assumes that ATP bound to any F-actin subunit throughout the filament hydrolyzes at the same rate. Indirectly, this implies that the conformation of the neighboring subunits does not significantly affect the rate of hydrolysis as the neighboring nucleotide state modulates the local conformational sampling of each monomer. Alternatively, the nucleotide state of neighboring subunits could modulate the rate, as is assumed in the cooperative hydrolysis models. Amongst the cooperative models, the vectorial model is in most contrast to the random hydrolysis model, since it assumes that ATP hydrolysis can occur predominantly in those subunits in the filament that have an adjoining ADP subunit, and hence are at an ATP-ADP (or ATP-ADP-P_i_) interface. In the strictest version of the vectorial model (32, 40), at the most two interfaces (exactly two if the filament is growing from both its ends) can exist in a filament, since hydrolysis is assumed to occur exclusively at the ATP-ADP interface and the interface simply moves towards the growing end of the filament as time progresses. The more realistic version of the vectorial model assigns a relatively small non-zero rate for ATP subunits that are not present at the interface (8).

There have been a number of studies attempting to perform a systematic comparison between the two classes of models (8, 32, 39, 41–43). However, common experimentally measured quantities such as rate of filament elongation, fluctuations in filament length and size of un-hydrolyzed ATP cap near the filament end are found to be insensitive to the mechanism of hydrolysis over a wide range of G-actin concentration, with small quantitative difference very close to the critical concentration (32, 43). A mixture of ATP bound G-actin and ADP bound G-actin can be used to introduce a different number of ATP-ADP interfaces in the filament by varying the composition of the mixture, thereby providing a way of enhancing the effective rate of hydrolysis within the context of the vectorial model (39). Fitting the predictions of a cooperative model to the time course of polymerization and ATP hydrolysis measured in these experiments eliminates the possibility of the strict vectorial model being accurate in all cases and suggests that the rate of hydrolysis at the ATP-ADP interface must be less than 100 times faster than the rate of hydrolysis away from the interface, although the predictions of a random hydrolysis mechanism were also shown to be able to explain the observed experimental data (39, 42). The random and vectorial models are based on two simple microscopic physical hypotheses that are able to accurately explain the experimental observations. An intermediate model could be constructed by assuming a more complex cooperativity that goes beyond the two-body binary cooperativity (two possible hydrolysis rates, one at the interface and one away from the interface) previously used in the vectorial model. However, lack of any direct experimental evidence differentiating these hypotheses makes empirically justifying and parameterizing such a complex cooperative model difficult.

As an alternative, we propose that one can develop models derived from all-atom (AA) molecular dynamics simulations that can provide a complex but detailed physical picture of this process based on the behavior produced by general atomistic force-fields. We constructed a UCG model that serves as a good representation of an actin filament at a low resolution, and also, for the first time, one that takes into account the nucleotide state of each actin subunit explicitly. The UCG filament model and conformational coupling between subunits was derived via a systematic procedure from AA molecular dynamics simulation, while the instantaneous conformation dependent rates of state transitions were approximated based on physical principles and implemented via a stochastic state hopping procedure with resulting macroscopic rates tied to experimental observations. After ensuring that the UCG model provided reasonable predictions for the mechanical properties of the filament, an MSM was constructed at the coarse resolution of traditional biological models. In the MSM, only the nucleotide composition along the position of subunits in the filament was retained, while conformational fluctuations were integrated out. This model was used to infer the spatial dependence of hydrolysis in actin filaments, allowing us to directly compare with established hydrolysis models in the literature. Finally, the MSM was then extended using experimental knowledge to explore the effects of concurrent hydrolysis and polymerization.

## COARSE-GRAINED MODEL

Our modeling approach is fundamentally based on using AA molecular dynamics simulations as the primary basis for constructing lower resolution CG models. The configurational behavior of the CG model of the filament conditional on nucleotide composition was parameterized in two steps, (1) using a systematic map to reduce the all-atom structure to fewer CG sites or “beads” (see Figure 1), and (2) using mapped system distributions to construct a CG effective force-field that governs the conformations of those coarse-grained beads in such a way that the all-atom behavior is faithfully reproduced. An essential feature of the UCG model in the current context is then the assignment of additional discrete internal states to the CG beads, with a different CG force-field associated with each state. In actin, for example, we used this additional model flexibility to represent the states of the bound nucleotide. The CG F-actin filament dynamics were then modeled using continuous time Langevin dynamics simulations of the CG beads using appropriate force-fields based on the instantaneous set of internal states of the neighboring CG beads, in addition to allowing discrete jumps between the various internal states. In other words, kinetics of ATP hydrolysis or P_i_ release in a subunit are controlled by the rates governing the switching between these states, which inherently must also take into account the conformational changes that the subunit undergoes upon hydrolysis or P_i_ release, and additionally taking into account the interdependence on the nucleotide states of its neighboring subunits. It is important to note that the latter significantly increases the complexity of the problem, and in order to make progress we must invoke certain assumptions. Firstly, we restricted the CG force-field to pair-wise interactions, and imposed a degree of locality on these interactions. As described in Ref. (33), without the locality approximation, there are exponentially large number of possible states to consider, making the UCG approach infeasible. The local nature of the interactions is somewhat based on biological intuition. Second, we constructed the CG force-field from AA simulations only for actin filaments consisting of subunits with identical nucleotides, and used a simple mixing rule in order to construct the CG force-field otherwise. A justification for the mixing rule is provided in the Supporting Material. The AA simulations, the UCG force-field, and the UCG discrete state-switching algorithm are described below.

**Figure 1:**
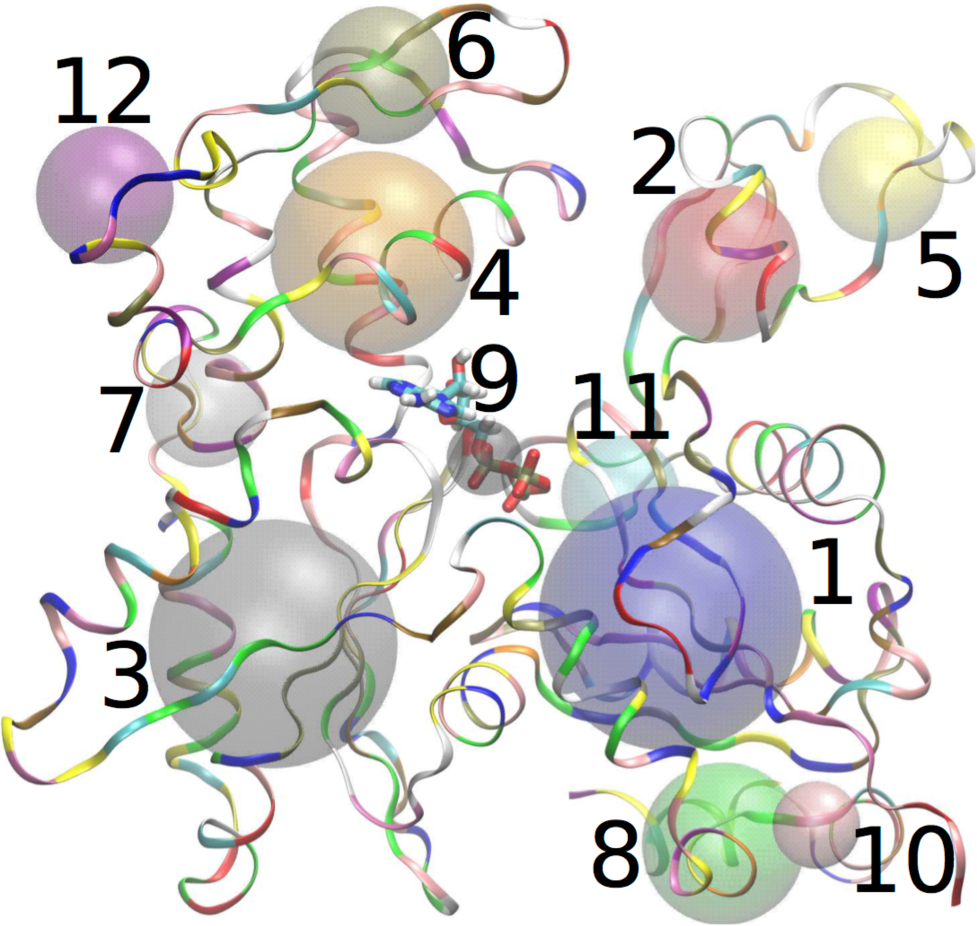
Schematic showing the CG mapping used in the present study. The atomistic structure of ADP-P_i_ bound actin subunit is shown as ribbons, with the corresponding CG sites shown as beads. CG bead indices 1 to 12 are marked next to each CG site. Our final model has 5 major beads correspond to CG sites with indices 1 to 5.

**Figure 2:**
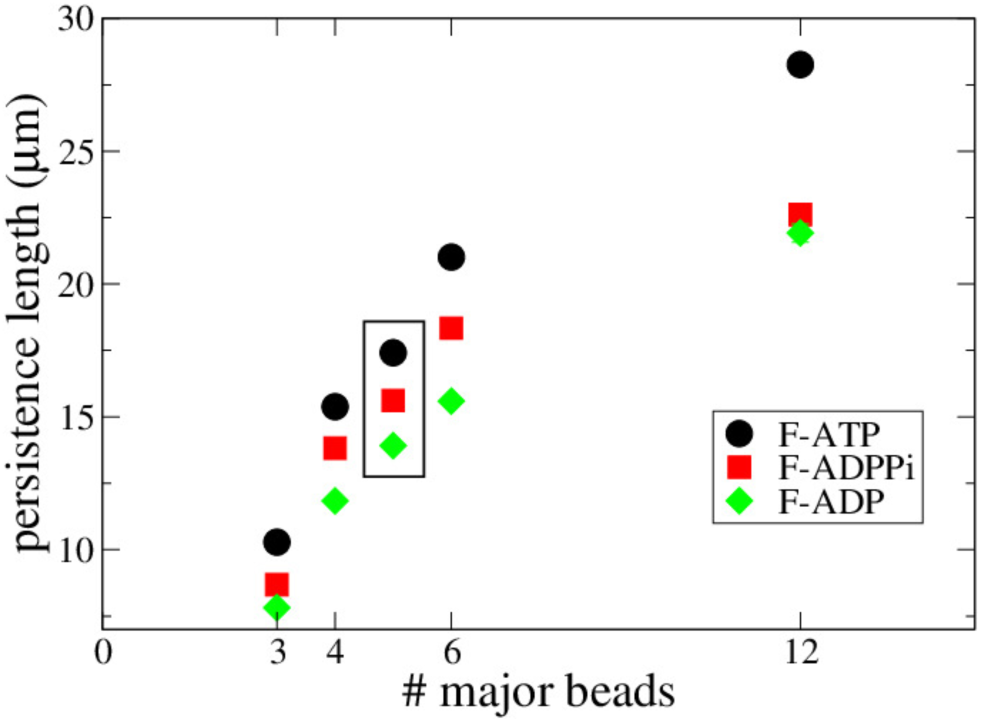
Persistence length of ATP, ADP-P_i_ and ADP bound actin filaments, as a funtion of the number of major beads. The major beads are chosen in the order of increasing CG indices.

### All-atom simulations

All-atom simulations of periodic actin filaments were performed in Gromacs (version 5.1.4)(44), using a protocol similar to Ref. (26). Briefly, a single actin subunit based on the Oda structure (Protein Data Bank structure 2ZWH) (45), consisting of a particular state of the bound nucleotide (either ATP, ADP-P_i_ or ADP, the former two were obtained by replacing the nucleotide ADP in the Oda structure with existing equilibrated simulations of ATP and ADP-P_i_ bound actin, with positions of replaced nucleotides found by aligning positions of actin subunits between the two (46)) was repeated along with a shift of 27.59 Å and a rotation of 166.6°, such that 13 repetitions of the subunit formed a single semi-periodic repeat of the actin helical structure. The simulation box was chosen such that the filament interacted with its own image in the periodic z-direction, mimicking a virtually infinite length filament. The system was solvated with TIP3P water and neutralized using salt ions, both of these tasks performed using VMD (47). The energy of the system was minimized, followed by gradual heating to increase the temperature from 0 K to 310 K, using the molecular dynamics code NAMD (48). The system was equilibrated at a temperature of 310 K and a pressure of 1 atm, until the root-mean-square deviation of the entire filament from its initial configuration reached a plateau. Production runs were performed using the terminal state of the system during equilibration, using the v-rescale thermostat (49) and the Parrinello-Rahman barostat implemented in Gromacs.

Three AA simulation trajectories were obtained, one for a pure ATP bound actin filament, one for a pure ADP-P_i_ bound actin filament, and one for a pure ADP bound actin filament. Each of these AA trajectories was used to obtain CG models for the filaments with corresponding states of the bound nucleotide.

### Ultra-Coarse-Grained force-field

Each of the ATP, ADP-P_i_ and ADP bound states of actin filament were independently coarsegrained using a 12-site mapping (26, 50) (Figure 1) and a hybrid force-field. The hybrid force-field consisted of an intra-subunit heterogeneous elastic network model (hENM, (24)), and an inter-subunit pair-wise interaction modeled with an inverted Gaussian potential, which will allow for future studies to include (de)polymerization. The details of these potentials are provided in the Supporting Material. First, AA simulation trajectories of an actin filament with 13 subunits, (bound exclusively with either ATP, ADP-P_i_ or ADP) were used to generate a force-field for a filament in a pure nucleotide state. To build the hybrid model, we started by connecting all pairs of intrasubunit beads by springs. We then connected inter-subunit beads with springs, but to simplify the conversion of inter-subunit interactions to be dissociable, we chose to only include springs between a subset of CG sites that we call *major beads* and restricted these inter-subunit springs to bead pairs that are less than three actin subunits apart to help enforce the locality of UCG interactions discussed earlier. Finally, we used the hENM procedure to assign spring constants that maximally match the fluctuations in the all-atom trajectory. To choose what CG sites to use for inter-subunit interactions, we tested varying numbers by of major beads, adding them in increasing order of their CG index as labeled in Figure 1. As discussed below, a model including the first 5 beads as major beads gave the best results and was chosen as our final model.

After the hENM procedure, the inter-subunit springs between actin monomers in the filament were converted into soft potentials. This was done by converting each inter-subunit elastic spring potential into an inverted Gaussian potential by least-square fitting to the elastic spring potential well in the region corresponding to a well depth of 3 kcal/mol. The well depth was set to be strong enough to prevent adjacent subunits in a filament from leaving the filament, while weak enough to prevent any large jumps in energy of the system that would lead to large numerical integration errors.

Figure 2 shows the variation in persistence length for the three states of bound nucleotide, obtained from the coarse-grained model simulated in LAMMPS MD software (51) using Langevin dynamics. Each data-point was calculated as an average over five simulation runs, with each initiated using a different seed for random force and initial velocity generation. For a given state of bound nucleotide, the filament became more flexible as the number of major beads decreased. Since the persistence length with 5 major beads (corresponding to CG bead indices 1 to 5 in Figure 1) agreed best with known persistence lengths in the literature, 5 major beads were used in the rest of the manuscript. Given the difficulty in accurately measuring persistence length in experiments and the wide range of experimental values reported in literature that varies with solution conditions, we picked a value that is consistent with the reported range (22, 24, 25, 52, 53). The 5 major beads in an actin subunit roughly correspond to the four major sub-domains in actin and the D-loop region (Figure 1). The D-loop region inserts into actin’s barbed end “target binding cleft” and is an important mediator of longitudinal interactions in the filament (20, 46, 53–57). This provided additional motivation for including at least these 5 major beads in our model.

The inter-subunit inverted Gaussian potential for a pair of CG beads with distinct states (say ATP:ADP-P_i_) of the bound nucleotide was obtained using a simple mixing rule that involved averaging parameters of the potential for each of the individual pure states (ATP:ATP and ADP-P_i_:ADP-P_i_), as described in the Supporting Material.

### Ultra-Coarse-Grained state switching

Each of the actin subunits was assigned an internal state depending on its bound nucleotide. The ATP hydrolysis and P_i_ release reactions were then represented as switching of these internal states (34). The simulation methodology consisted of evolving continuous variables (positions, velocities of coarse-grained beads) using Langevin dynamics, along with discrete state transitions. Physically, these discrete transitions correspond to either hydrolysis or phosphate release, with the instantaneous rate an expression of how the barrier of the reaction changes depending on the instantaneous configuration of the filament. The nature of this dependence indirectly produces all hydrolysis and phosphate release cooperativity observed in the current study.

However, full rigorous parametrization of this dependence, either through experimental data or reactive atomistic simulation, is infeasible. Experimental data cannot achieve the required resolution, and a reactive atomistic simulation is computationally prohibitive when considering the dependence of a complex reaction on a multitude of protein environments. Instead, simple arguments on the transition state stability as a function of the reactants or product stability were used. Fundamentally, the approach is similar to kinetic implications of the Hammond postulate: the free energy of the transition state has approximately the same dependence on configuration as either the products or reactants, depending on which of the products or reactants is closer in free energy to the transition state (58). Additional dependence of the transition state free energy on the local configuration, such as the dihedral angle of the monomeric unit, was introduced through terms *k*(*ϕ*) in the equations below. This approach resulted in the following instantaneous rate expressions. For a given subunit initially in the state *i*, the instantaneous rate of switching to state *j* is given by the following equation based on the Metropolis-Hastings-like criterion

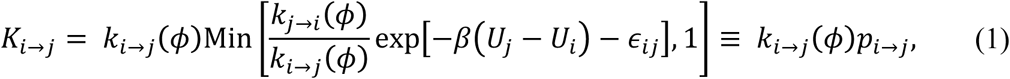

where

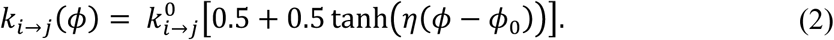

Here, *U_j_ − U_i_* is the energy difference between states *j* and *i, ϕ* − *ϕ*_0_ is the difference between the instantaneous dihedral angle between CG beads 2-1-3-4 of the subunit and its average value in state *i, η* is the parameter that controls the explicit dihedral angle dependence, while 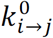, and ∊*_ij_*, are model parameters. Computationally, the prefactor *k_i→j_(ϕ)* can be seen as specifying the rate of attempting a state transition, and *p_i→j_* gives the probability of accepting the transition. As the transition describes a chemical reaction in an equilibrium system, detailed balance implies that the rate for the reverse reaction, in which a subunit initially in state *j* switches to state *i*, is given by

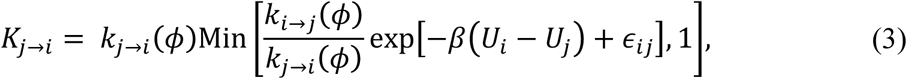

where

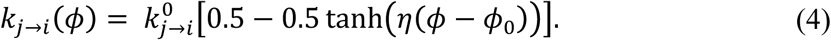

The explicit dihedral angle dependence was modeled as a smooth step function such that it resulted into an increase in *k_i→j_(ϕ)* for the forward reaction and decrease in *k_j→i_(ϕ)* for the reverse reaction, as the dihedral angle *ϕ* increases. The explicit dihedral angle dependence was based on our previous work (4–6) that attributes the increase in rate of hydrolysis to the flattening of the actin subunit. We set *ϕ*_0_ = −10° and used *η* = 0.125 for the ATP hydrolysis reaction, but turned off the explicit dihedral angle dependence for the P_i_ release reaction by setting *η* = 0. Additional discussion of the physical meaning of these parameters can be found in the Supporting Material.

### Parameter estimation

The parameters 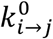, 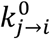 and *∊_ij_*, are the UCG model parameters that need to be estimated. These were optimized using the following three conditions. The instantaneous forward and reverse reaction rates, on an average, must match the known macroscopic reaction rates. This provides two conditions, one for the forward reaction and one for the reverse reaction. These two conditions were specified by the system being simulated. The third condition has more freedom in its choice, and we selected this condition such that the average acceptance probability for the forward reaction had a desired value. The motivation behind this choice was that it provides a handle to control the sensitivity of state transitions to the energy difference *(U_i_ − U_j_)* between the two states. The details of parameter estimation are provided in the Supporting Material (5, 6, 59, 60). In the final model, we set the average acceptance probability to 0.1 for both ATP hydrolysis and P_i_ release reactions.

### Markov state model

The UCG simulations performed were analyzed through their behavior at the resolution of an MSM. The MSM resolution was based on the kinetic models used in the literature (31, 43). In this MSM description, an actin filament contains no configurational behavior; we only considered the length of actin filaments, and their composition in terms of the nucleotide state. In other words, the actin filament system was represented by a state vector, with positions of elements of the vector corresponding to positions of actin subunits in a filament and the value of each element (ATP, ADP-P_i_ or ADP) corresponding to the state of the nucleotide bound to the respective subunit. Mean first passage times were estimated using the mapped statistics observed in the UCG simulations and were used to parameterize the MSM. This procedure is described in more detail in the discussion section.

The MSM model was further extended to include polymerization and depolymerization of actin units. As the addition of ATP bound actin to the filament introduces energy into the local system represented by the MSM, the constraint of detailed balance was not imposed in when considering the transitions related to the addition or removal of actin units. As a result, when considering polymerization, the length of the state vector at a given instant was equal to the length of the filament at that instant, and the two terminal positions of the state vector corresponded to the barbed and pointed ends of the filament. Polymerization (de-polymerization) at the ends resulted in expansion (shrinking) of the state vector, while ATP hydrolysis and P_i_ release of a particular subunit resulted in a change in the value of the corresponding element of the state vector. To further understand the importance of the cooperativity in the filament, the rate parameters via the results of the UCG model were uniformly scaled to probe the effect of increased or decreased cooperativity. The model was sampled using a Monte Carlo algorithm (see Supporting Material for details).

## MULTI-BODY EFFECTS IN KINETICS OF ATP HYDROLYSIS AND P_I_ RELEASE

The UCG model was used to study multi-body effects in ATP hydrolysis and P_i_ release at the resolution of the MSM model. The hydrolysis of each actin subunit can be affected by several neighboring subunits. Ideally, one needs to consider all possible combinations of the nucleotide states of several neighboring units in order to study their effect on the rate of hydrolysis. To keep the number of such combinations tractable, we limited the study to a fairly small number of neighboring subunits by invoking the local nature of interactions between coarse-grained beads. The longest pair-wise interactions in our model were between coarse-grained beads belonging to actin subunits that are two monomers apart in the filament. Hence, we limited our study of multibody effect to three neighboring subunits on each of the two sides of a given subunit, as described below.

For ATP hydrolysis, we designed a long UCG filament model consisting of 18265 actin subunits as follows: We capped the barbed end of the filament with 26 ATP bound subunits and the pointed end with 39 ATP bound subunits to avoid any possible end-effects. The remaining 18200 subunits in the “bulk” of the filament were divided into sets of 7 subunits long sections. The 4^th^ subunit (marked subunit) in each section was an ATP bound subunit, while three of its nearest neighbors towards the pointed end (1–3) and towards the barbed end (5–7) were randomly chosen to be either ATP bound or ADP-P_i_ bound subunits. In practice, the nucleotide states of the six neighbors of a marked subunit in each of the 2600 sections were randomly chosen from all 2^6^ = 64 possible combinations, such that there were at least 40 copies of each combination in a single filament at random locations along its length. Only marked subunits were allowed to hydrolyze, while all the neighbors simply underwent Langevin dynamics, and were constrained to remain in their initial nucleotide state. UCG simulations were run until all marked subunits hydrolyzed. These simulations were repeated 640 times, each with a unique filament design, and the conditional mean first passage time (MFPT) for each combination of neighboring states was calculated. The inverse of the mean first passage time for all the hydrolysis events provides the rate for that reaction in the UCG parametrized MSM when considering reactions at the full granularity of neighbors (61). At coarser resolutions, we refer to the corresponding effective rate as the average rate, given by

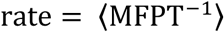

In the following, a unique six-digit *key* is used to denote a section with a particular combination of neighboring subunits. An un-hydrolyzed ATP bound subunit is represented as 0, while a hydrolyzed ADP-P_i_ bound subunit is represented as 1. The key is simply the word formed by concatenating these representations, starting with the third nearest neighbor towards the pointed end and going through each consecutive neighbor up to the third nearest neighbor towards the barbed end.

Figure 3 (and Figure S4(a) in Supporting Material) shows the variation in rate of ATP hydrolysis relative to the average rate as a function of certain representative combinations of states of neighboring subunits. The explicit dihedral angle dependence in Equations (2) and (4) was used, with *η* = 0.125. The ATP hydrolysis rate of a given subunit was found to vary by a maximum of about +/− 20% relative to its average rate, depending on whether it is in an ATP rich or an ADP-P_i_ rich environment. The reaction rate was enhanced when all six neighboring subunits were in the un-hydrolyzed ATP bound state, while it was suppressed when all these neighbors were in the ADP-P_i_ bound state. Similar variation in hydrolysis rate was observed even when ignoring the dependence of the two farthest neighbors (data corresponding to keys X0000X and X1111X in Figure S4(a) in Supporting Material), supporting our assumption of the local nature of interactions. The observed variation was found to be insensitive to the explicit dihedral dependence (Figure S4(b) in Supporting Material).

**Figure 3:**
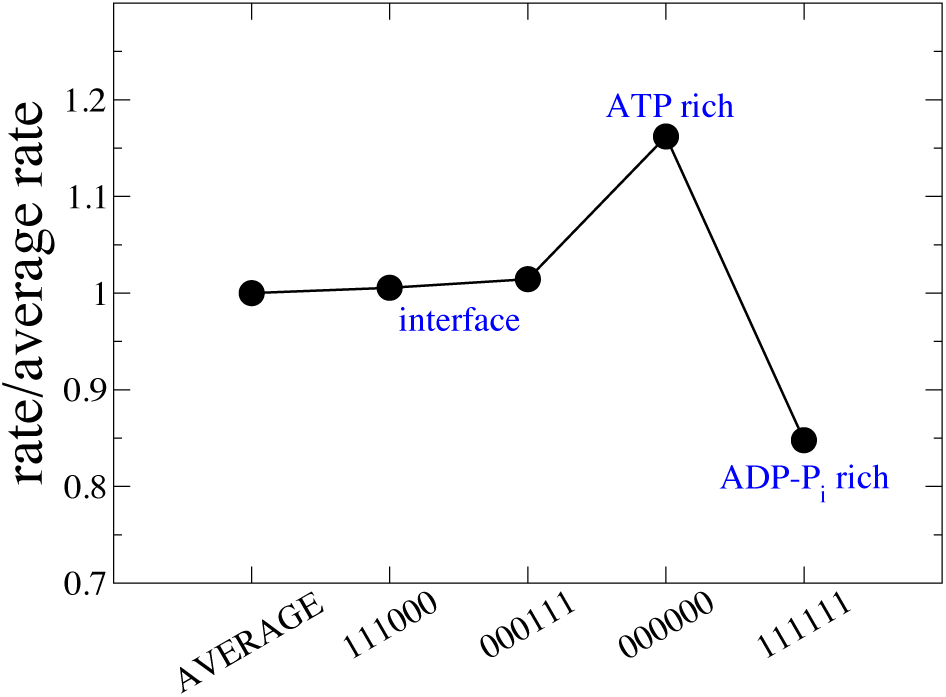
Multi-body effect in ATP hydrolysis, plotted as a ratio of the conditional rate with the average rate for specific combinations of nucleotide states of neighboring subunits. Combinations of neighboring subunit states are indicated using a key on the x-axis, that denotes the state (0=unhydrolyzed, 1=hydrolyzed) of each of the neighboring subunits, starting from the third neighbor towards the pointed end to the third neighbor towards the barbed end. Error bars indicate the standard error for each data-point, and are smaller than the symbol size for most of the data. See Figure S4(b) in Supporting Material for data corresponding to all 64 possible combinations.

For P_i_ release, we designed long actin filaments consisting of 18265 actin subunits, similar to those used in the above simulations. The barbed and pointed ends of the filaments were capped with ADP-P_i_ bound subunits, and the “bulk” of the filament was divided into 7 subunit long sections. Each section was randomly chosen such that the 4^th^ subunit was initially ADP-P_i_ bound and was allowed to release its P_i_, while the rest of the subunits mimicked all 64 possible combinations of states (either ADP-P_i_ or ADP bound) and were forced to remain in their initial state. Again, large statistical sampling was used in obtaining the kinetic data, where the mean first passage time for each particular combination of neighbors was obtained from 640 filaments, each containing at least 40 copies of the combination at random locations along the filament.

The P_i_ release reaction rate was also affected by the state of neighboring subunits by a maximum of about +/− 20% over its average rate, as seen from Figure 4 (and Figure S6 in Supporting Material). Note that the explicit dihedral angle dependence in Equations (2) and (4) was switched off in the P_i_ release reaction by setting *η* = 0. In contrast to ATP hydrolysis, the P_i_ release rate of a given subunit was enhanced when all of its neighbors were free of P_i_. The rate was suppressed when all of its neighboring subunits were ADP-P_i_ bound. At the interface, where all three neighbors along the pointed end side of the marked subunit were ADP bound and those along the barbed end side of the marked subunit were ADP-P_i_ bound (corresponding to key 111000), the rate of reaction was only marginally higher relative to its average value. The observed variation was found to persist within a range of the chosen value for average acceptance probability (see Figure S5 in Supporting Material).

**Figure 4:**
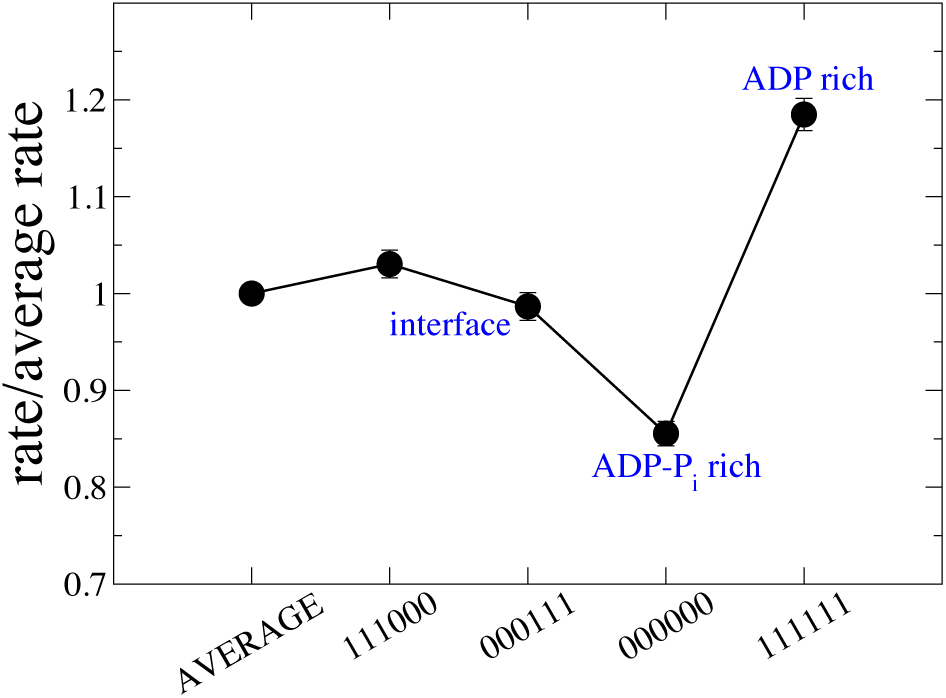
Multi-body effect in P_i_ release, plotted as a ratio of the conditional rate to the average rate for specific combinations of nucleotide states of neighboring subunits. The key on the x-axis is similar to that described in Figure 3 (except for the new definitions 0=ADP-P_i_, 1=ADP). Error bars indicate the standard error for each data-point. See Figure S6 in Supporting Material for data corresponding to all 64 possible combinations.

## FILAMENT GROWTH AND ROLE OF MULTI-BODY EFFECTS

The role of ATP hydrolysis and P_i_ release on filament growth has been previously studied using kinetic models that include polymerization and de-polymerization of subunits at the ends of the filament, along with ATP hydrolysis in subunits belonging to the filament (31, 32, 43). These models have been shown to agree well with experimental data in terms of the average filament growth rate, independent of whether the vectorial (32) or stochastic (31) mechanism of hydrolysis is assumed. In these models, multi-body effects were ignored to keep the models tractable and possibly due to lack of any evidence suggesting such effects. On the other hand, our UCG model predicted a strong multi-body effect, manifested through a significant variation in both the ATP hydrolysis and the P_i_ release kinetics depending on the state of neighboring subunits. In the following, we studied the implications of this multi-body effect on filament kinetics and statistics using our MSM based on kinetic models similar to Refs. (31, 43). This is performed by extending the previous MSM characterizing only hydrolysis via the addition of states and transitions representing polymerization and depolymerization. As described below, the primary distinguishing feature of our MSM was the explicit dependence of rates of ATP hydrolysis and P_i_ release on the nucleotide states of neighboring subunits, based on the predictions of our UCG model (Figure 3, Figure 4 and Figure S4(b), Figure S6 in Supporting Material).

The filament was initialized with *n*_0_ subunits and was assumed to be in a solution of free monomeric subunits at a specified concentration. Addition of a free ATP bound subunit at the barbed end of the filament increased the filament length, while dissociation of the terminal subunit (ATP or ADP-P_i_ or ADP bound) at the barbed or pointed ends decreased its length. The rest of the subunits belonging to the filament underwent ATP hydrolysis and P_i_ release. The macroscopic rate constants for these reactions are given in Table 1, and were chosen to be same as in the original models with which we compared our predictions.

**Table 1:** Rate constants used in the kinetic model (31, 43). K_pol_. is the polymerization rate. K_dis_., _barbed_ is the de-polymerization rate at the barbed end, while K_dis_., _pointed_ is that at the pointed end. K_hyd_. and K_rel_. are rates of ATP hydrolysis and P_i_ release respectively.

| $K_{pol.} (\mu M^{-1} s^{-1})$ | | $K_{dis., barbed} (s^{-1})$ | | | $K_{dis., pointed} (s^{-1})$ | | $K_{hyd.} (s^{-1})$ | $K_{rel.} (s^{-1})$ |
| --- | --- | --- | --- | --- | --- | --- | --- | --- |
| barbed | pointed | ATP | ADP-$P_i$ | ADP | ATP | ADP | | |
| 11.6 | - | 1.4 | 1.1 (31) | 7.2 | 0.8 (43) | 0.27 (43) | 0.3 (31) | 0.003(43),<br>0.004(31) |

The model was sampled using a Monte Carlo (MC) algorithm to study evolution of the filament, keeping track of the location of all the actin subunits in the filament. For each set of parameters, 1000 statistically independent simulations were run. Details of the MC algorithm are provided in the Supporting Material.

### Filament dynamics in a conserved system

A conserved system (where the total number of subunits in the system, including free monomers and polymerized subunits, remains constant in time) was proposed in Ref. (43) to study the effect of vectorial versus stochastic hydrolysis on the transient part (see Figure 5) of filament growth. For the conserved system, starting with an initial filament of a certain length and an initial free monomer concentration c_0_ that is above the critical concentration at which the net filament growth rate is positive, the free monomer concentration keeps on dropping as the filament grows, thereby reducing the polymerization rate. This continues until the concentration is just enough to balance the net polymerization and dissociation rates, resulting into a nearly steady filament length at larger times. It was predicted that although the vectorial and random mechanisms would result into a similar steady filament length, the transient from the initial filament to the steady filament would show significant variation depending on the mechanism of hydrolysis. For a direct comparison with Ref. (43), we further modified our model such that the phenomenology between the two models was exactly the same. Firstly, ATP hydrolysis was assumed to be fast relative to P_i_ release. Hence, ATP and ADP-P_i_ were made indistinguishable in the model. Further, the filament was allowed to polymerize only at the barbed end. The resulting model consisted of only two states of the nucleotide bound to actin subunits in the filament: ATP and ADP. The rate constants in Table 1 corresponding to Ref. (43) were used.

**Figure 5:**
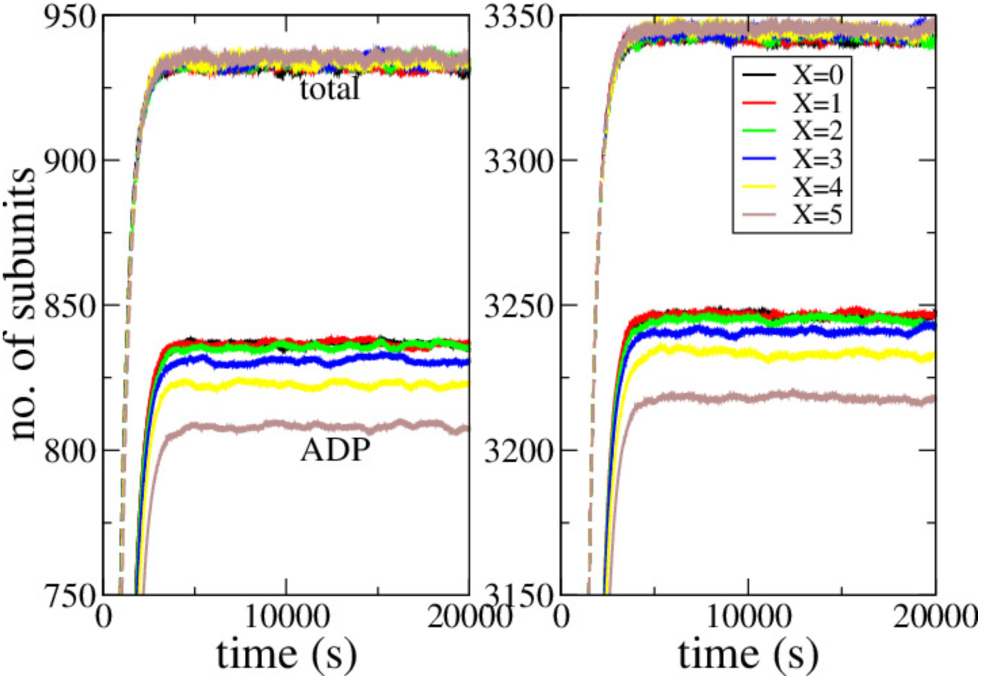
The total number of subunits (dotted lines) and the number of hydrolyzed subunits (solid lines) in the filament, as a function of simulation time. Different colors correspond to varying strength of multi-body effects incorporated in the model, with X = 0 corresponding to a purely stochastic hydrolysis. Left panel corresponds to c_0_=0.3 μM, while the right panel corresponds to c_0_=0.7 μM.

In order to incorporate the multi-body effects as revealed in the UCG simulations into the model, an additional modification was made. As already mentioned, our Monte Carlo algorithm was designed to keep track of the locations of all actin subunits in the filament. This allowed us to use all of the data in Figure S6 (in Supporting Material), since we knew the instantaneous nucleotide state of neighbors of each actin subunit. We introduced a parameter *X* in the model, as discussed below, to have the ability to interpolate between a purely stochastic mechanism (*X* = 0) and the UCG predictions (*X* = 1). Since the multi-body effects predicted by our UCG model were sensitive to the UCG parameters (although similar in trends, see Figure S5 in Supporting Material), the parameter *X* also allowed us to extrapolate beyond the UCG model predictions specific to the choice of UCG parameters in our final model. The multi-body rate of P_i_ release, 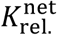, was defined as follows in order to take into account the variation with state of neighbors:

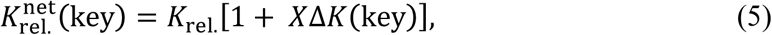

where

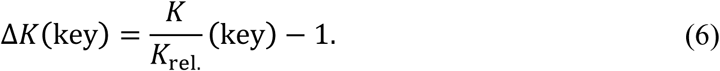

The first term of the right-hand side of Equation (6) is simply the ratio of conditional rate of P_i_ release *K* for a combination of states of neighboring subunits specified with the key, to the average rate of P_i_ release *K*_rel_ for all such possible combinations (same quantity as the y-axis in Figure 4 and Figure S6 in Supporting Material). For example, the ratio 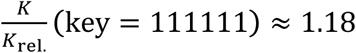 for a subunit surrounded by all ADP bound neighbors, as observed from Figure 4. The rate of P_i_ release for each ATP bound subunit was similarly modulated by using in Equation (6) the ratio 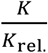 from Figure S6 corresponding to its instantaneous key at each Monte Carlo step. When a ATP bound subunit released its P_i_ during a Monte Carlo step with a P_i_ release rate governed by the instantaneous nucleotide state of its neighbors, the keys of all its neighboring sub-units were updated at the end of the step, which in turn modulated the P_i_ release rates of its neighboring ATP bound subunits in all future Monte Carlo steps. As mentioned earlier, the parameter *X* was used to vary the strength of multi-body effects in the model. The specific values of UCG parameters chosen in this work corresponded to *X = 1*. However, acknowledging the possibility of other choices for the UCG parameters, we allowed *X* to vary up to *X = 5* (since 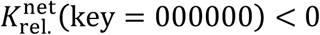 for *X* = 6 and beyond).

Figure 5 shows the multi-body effect on mean filament dynamics observed across 1000 simulation runs, for two values of initial concentration *c_0_*. The number of hydrolyzed subunits, initially set to 4, underwent a transient, beyond which it remained nearly constant. Although the qualitative features were similar across all strengths of multi-body effects studied, there was an increasing delay in hydrolysis relative to the filament growth dynamics as this strength increased. Moreover, the asymptotic value of hydrolyzed subunits systematically decreased. The total number of subunits and hence the filament length remained nearly constant after undergoing a transient from its initial value of 6. However, both the transient and the asymptotic value of the filament length were not found to be sensitive to the strength of multi-body effects.

On the other hand, the filament composition showed a stronger dependence on the mechanism of P_i_ release. A contiguous ATP section, throughout which all the consecutive subunits were ATP bound, could easily be identified in our simulations. At every instant, we identified the number and length of these sections in the filament, and calculated their mean values across all 1000 simulation runs. Figure 6(a) shows the average length of a contiguous ATP section as a function of time. In comparison to a purely random P_i_ release, the multi-body cooperative effects predicted much longer contiguous ATP sections. The same trend was also observed in the mean of the maximum length of these sections across each simulation run, as shown in Figure 6(b).

**Figure 6:**
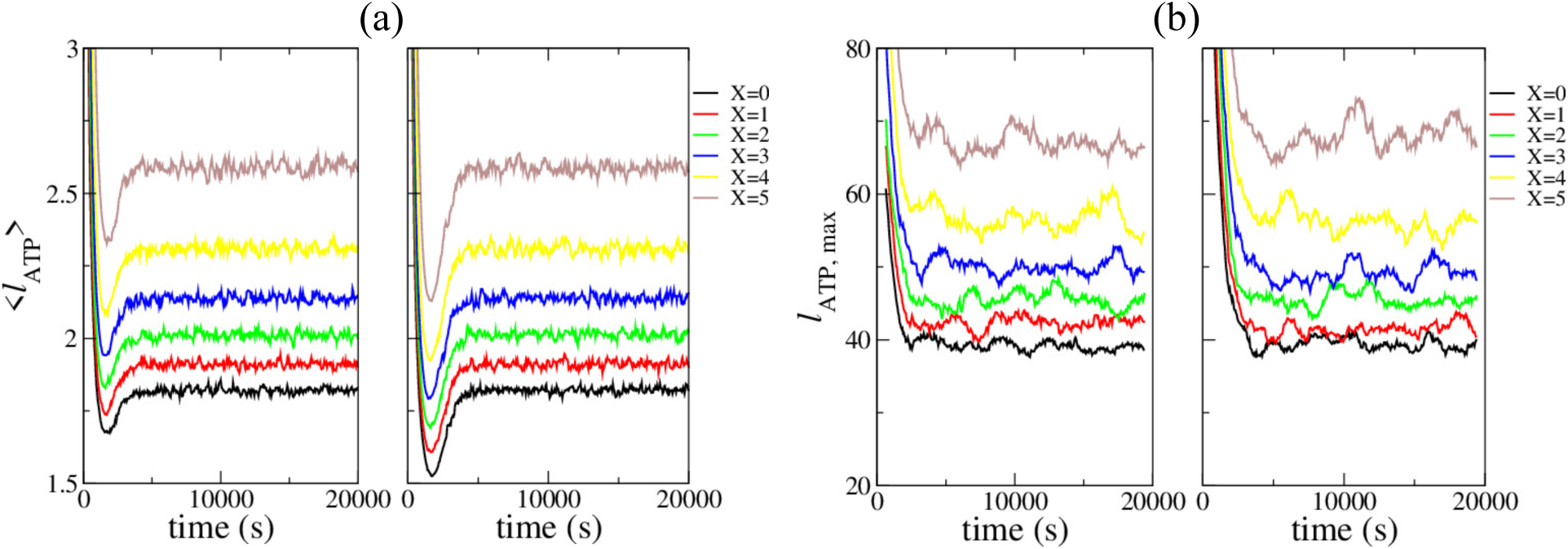
Variation in filament composition due to incorporation of multi-body effects in P_i_ release reaction, shown as (a) average length of a contiguous ATP section along the filament and (b) maximum length of a contiguous ATP section along the filament. Different colors indicate varying strength of multi-body effects, as indicated in the legend. Left panel corresponds to c*_0_*=0.3 μM, while the right panel corresponds to c_0_=0.7 μM.

### Filament dynamics at constant free actin concentration

An alternative to the aforementioned approach is to keep the total free actin concentration constant. This mimics experimental conditions where, for example, a solution of G-actin at constant concentration is flowing through the system using microfluidic devices. Since the total concentration of free actin does not change with time, the resulting filament either grows with time if the free actin concentration is above the critical concentration, or shrinks with time if the concentration is lower than the critical concentration.

The filament growth rates at different free actin concentrations obtained in experiments are shown to be in agreement with either of the vectorial or stochastic mechanisms of hydrolysis (31, 32, 43). We modified our MSM based on the kinetic model in Ref. (31), by using corresponding parameters from Table 1. For direct comparison with Ref. (31), we included all three possible bound nucleotide states of subunits in our model: ATP, ADP-P_i_ and ADP. Additionally, we assumed that no polymerization or dissociation took place at the pointed end. Thus, the model consisted of polymerization and dissociation at the barbed end, with ATP hydrolysis and P_i_ release throughout the filament. We further modified the model to incorporate the multi-body effects observed in our UCG simulations, by modulating the ATP hydrolysis and P_i_ release rates using the data in Figure 3 and Figure 4 (and Figure S4(b) and Figure S6 in Supporting Material) as follows. Given a state of neighboring subunits with a corresponding key, the net rate of ATP hydrolysis is defined as

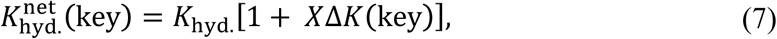

where

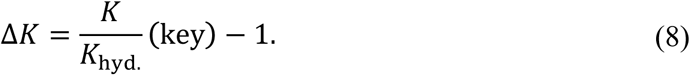

Here, the first term on the right-hand side of Equation (8) is the ratio of conditional rate of ATP hydrolysis to the average rate observed in our UCG model, plotted in Figure 3. The net rate of P_i_ release was obtained using Equations (7) and (8). The strength of multi-body effects was varied using the parameter *X*, with *X = 0* corresponding to a purely stochastic mechanism. As discussed earlier, our specific choice of UCG parameters corresponded to *X = 1*, and we treated *X* as a parameter to acknowledge the possibility of other choices for UCG parameters.

The free actin subunits were implicitly present at a constant concentration *c*. The filament initially consisted of *n*_0_ subunits, with 2/3^rd^ fraction of the filament near the pointed end being ADP bound, and the reminder tip near the barbed end made of ATP bound subunits.

Figure 7(a) shows the mean total filament length (number of subunits) obtained from 1000 statistical runs, for two different free actin concentrations. The left panel corresponds to a free actin concentration below the critical concentration. The filament, initially made of *n_0_* = 2000 subunits, shrunk at a constant rate as subunits dissociated. The right panel corresponds to a free actin concentration above the critical concentration. The filament, initially made of *n_0_* = 1000 subunits, grew at a constant rate. For both concentrations, the filament growth followed a different trajectory depending on the strength of multi-body effects incorporated into the model.

**Figure 7:**
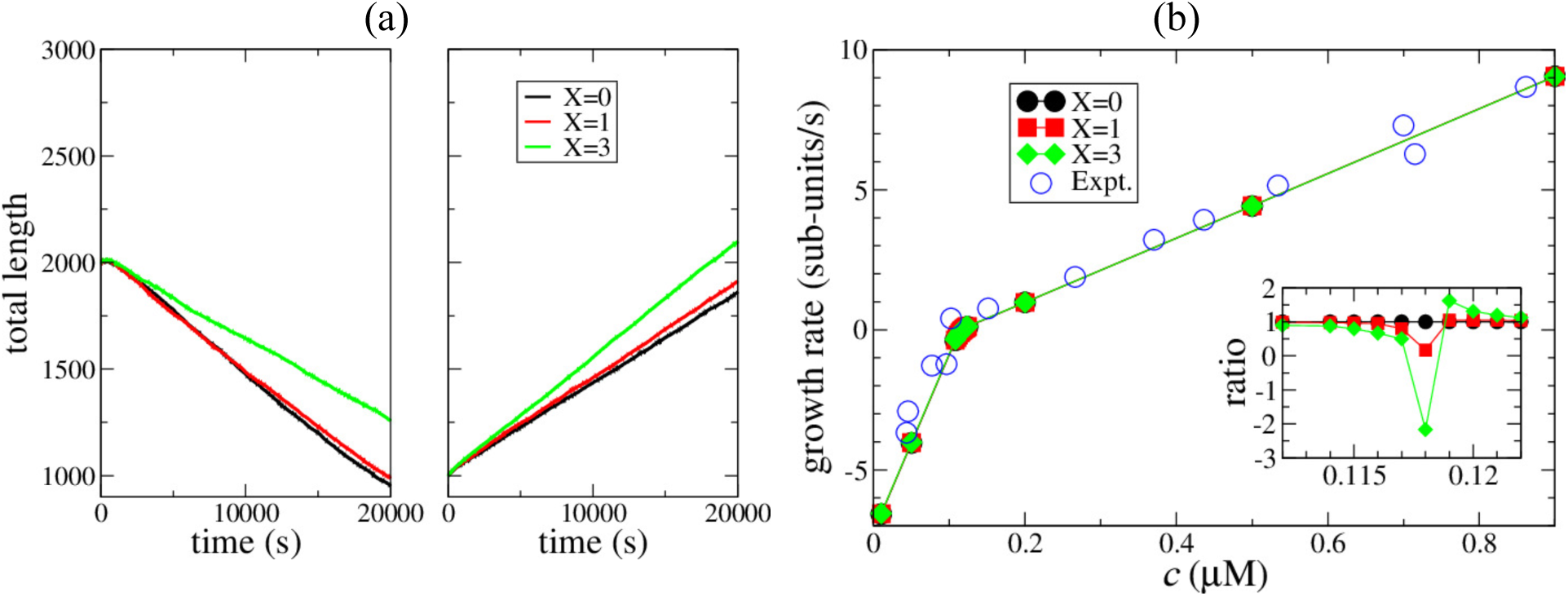
(a) Filament length dynamics for different strengths of multi-body effect shown in different color, at concentrations below (left panel, c = 0.16 μM) and above (right panel, c = 0.19 μM) the critical concentration. (b) Filament growth rate (filled symbols) as a function of concentration of free actin. Different colors represent different strengths of multi-body effects. Open circles are experimental data taken from Ref. (31), originally extracted from experiments in Ref. (43). Inset shows the ratio of growth rate at a given strength of multi-body effects compared to the growth rate at *X* = 0.

The slope of the filament length curves, ignoring the initial transient, gives the growth rate of the filament. Figure 7(b) shows the growth rate of the filament, as a function of the free actin concentration. It is evident that the filament growth rate is not dramatically affected by absence (*X* = 0) or presence (*X* ≠ 0) of the multi-body effects. Note that small differences in the growth rate can affect the filament length trajectories significantly, especially at longer times. Near the critical concentration, at which the growth rate is zero, the strength of multi-body effects incorporated in the model affected the growth rate significantly, as the inset of Figure 7(b) shows. However, the absolute growth rate near the critical concentration was too low to make these variations significant.

Although the filament growth kinetics were not affected significantly by the mechanism of hydrolysis and P_i_ release, the composition of the filaments changed significantly as multi-body effects were made stronger in the model. As the filament grew at a constant rate above the critical concentration, more and more ATP subunits were added to its barbed end. These subunits then underwent hydrolysis and P_i_ release. As the model ignored any dissociation at the pointed end, the number of ADP subunits also grew with time. On the other hand, the number of ADP-P_i_ subunits remained nearly constant after an initial transient. Figure 8(a) shows the average length of a contiguous ADP-P_i_ section for a range of free actin concentrations obtained as a mean over the nearly constant regime and over 1000 simulation runs. At a given free actin concentration, the average length increased with increasing strength of multi-body effects. This trend was also reflected in the mean value of maximum length of a contiguous ADP-P_i_ section shown in Figure 8(b).

**Figure 8:**
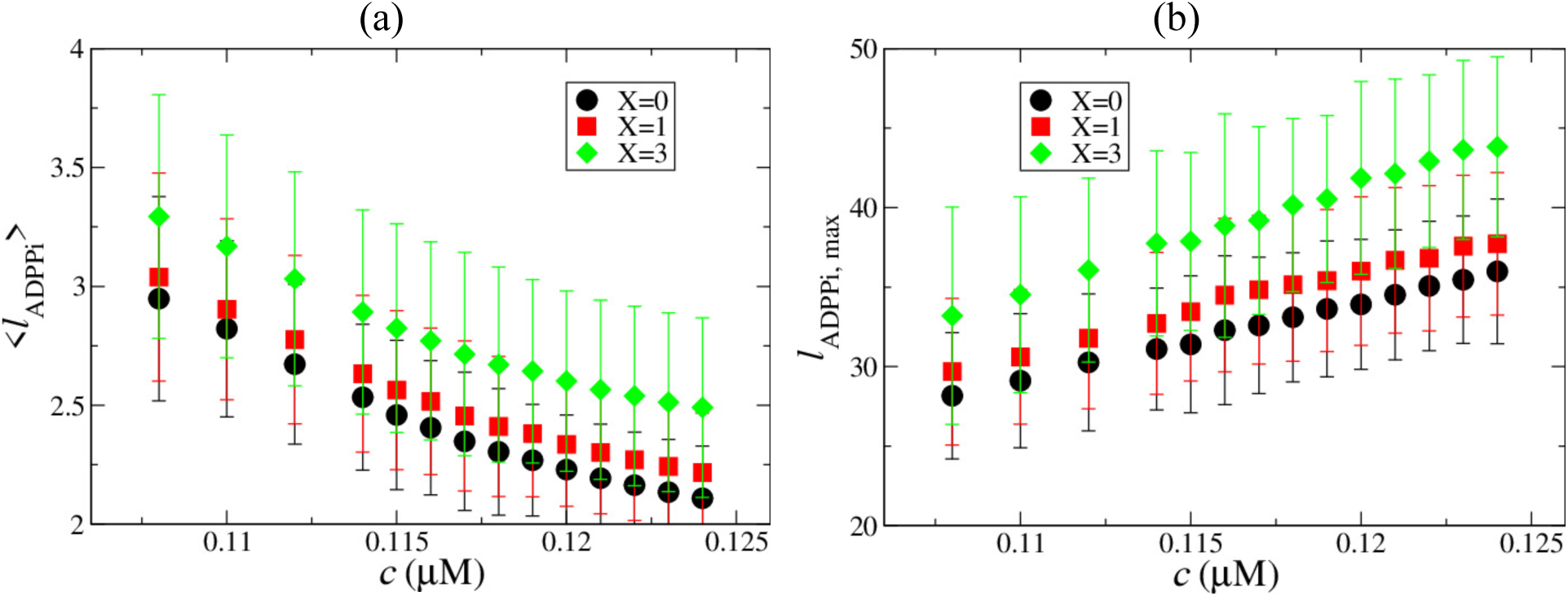
Variation in filament composition due to incorporation of multi-body effects in ATP hydrolysis and P_i_ release, shown in terms of (a) the average length of a contiguous ADP-P_i_ section along the filament, and (b) maximum length of a contiguous ADP-P_i_ section along the filament. Different symbols indicate varying strength of multi-body effects, as indicated in the legend.

## DISCUSSION

Two classes of kinetic models have been proposed for actin filaments, based on the mechanism of hydrolysis of ATP and the subsequent release of P_i_. The random hydrolysis mechanism (2, 31, 39, 42) assumes an equal probability of ATP hydrolysis and release P_i_ at every position in the filament. On the other extreme, the vectorial model (8, 32, 40, 41) assumes that hydrolysis can take place only at the boundary between the un-hydrolyzed and hydrolyzed parts of the filament. Although the two models fundamentally differ in terms of hydrolysis, the difference seems to be insignificant for filament dynamics, especially in terms of the filament growth rate that is typically measured in experiments.

The random hydrolysis mechanism assumes no cooperativity in hydrolysis, so that subunits at any position along the filament hydrolyze at the same rate at all times. The vectorial mechanism assumes maximal cooperativity in hydrolysis, such that only those subunits that are at the interface between un-hydrolyzed and hydrolyzed units undergo hydrolysis at a non-zero rate, and all other subunits throughout the filament effectively have a zero rate of hydrolysis. Our UCG and kinetic models allowed us to explore the possibility of intermediate levels of cooperativity, where the nearby neighbors affect the rate of hydrolysis of a given subunit in a filament.

More specifically, the UCG model predicted a substantial variation in the rates of hydrolysis and P_i_ release. The rate of ATP hydrolysis was enhanced by about 20% when an ATP actin subunit was in an ATP rich environment and decreases by about 20% when in an ADP-P_i_ rich environment. In the scenario of a growing actin filament, such a variation implies that the filament would be more fragmented in ATP in comparison to random hydrolysis, because subunits in the ATP rich region will hydrolyze faster to form ADP-P_i_, which then slows down the hydrolysis of nearby neighbors to a certain extent. Data from our kinetic model reflected this in terms of the average and maximum lengths of ATP sections (not shown). In contrast, the rate of P_i_ release from an ADP-P_i_ subunit was suppressed when it was in an ADP-P_i_ rich environment and enhanced by about ~20% when in an ADP rich environment. This implies that in a growing filament, the filament would be less fragmented in ADP-P_i_ in comparison to random hydrolysis, because subunits in the ADP-P_i_ rich region of the filament will release their P_i_ slower to form ADP, which will then accelerate the P_i_ release of neighboring subunits to a certain extent. This is exactly what we observed in our kinetic model, as shown by the data in Figure 8.

An important point to consider is that the hydrolysis reaction is faster than the P_i_ release reaction by about two orders of magnitude (Table 1). Although the relative variation in rates of hydrolysis and P_i_ release are of the same extent in Figure 3 and Figure 4, the variation in absolute rate of the faster hydrolysis reaction is expected to play an insignificant role in filament dynamics and composition. The slower P_i_ release reaction is the rate determining step, and hence should have the strongest effect on filament dynamics and composition.

Although the filament composition changes significantly with the cooperativity in ATP hydrolysis and P_i_ release, long length-scale filament properties such as the persistence length do not reflect this change to a measurable extent. As seen from Figure 5, in a conserved system, a steady state characterized by a filament length that fluctuates around a constant average value was reached as the actin monomer concentration approached the tread-milling concentration. For systematic comparison, we chose an initial actin concentration c_0_=0.2 that gave a filament of average length 325 subunits. We listed about 90 unique compositions that the filament of length 325 subunits exhibited, and calculated the average persistence length based on these compositions. The average persistence length of the filament under these conditions was found to be 13.5 (± 0.2) μm for the random mechanism corresponding to *X* = 0 cooperativity, 13.6 (± 0.2) μm for *X* = 3 cooperativity, and 13.9 (± 0.2) μm for *X* = 5 cooperativity. When the free actin concentration is kept constant, the filament is either growing or shrinking at a constant rate with time, except at the critical concentration for which the growth rate is zero. Since there is no steady state with constant filament length in this system (except at critical concentration), we listed 90 unique compositions for a filament length of 325 subunits based on MC runs of the system at c_0_=0.120 (just above the critical concentration, see Figure 7(b)) to calculate the persistence length. The average persistence length for a filament length of 325 subunits for the random mechanism corresponding to *X* = 0 cooperativity was found to be 14.2 (± 0.2) μm, while that for *X* = 3 cooperativity was found to be 14.4 (± 0.2) μm. The changes in persistence length corresponding to the changes in composition were expected to be small, given the narrow range between persistence lengths for a pure ADP-P_i_ filament and a pure ADP filament (Figure 2).

In conclusion, our bottom-up coarse-graining strategy enabled us to probe the cooperative nature of ATP hydrolysis and P_i_ release in F-actin in detail. We found that the mechanism of these reactions is not fully random, but does depend on the state of neighboring subunits. Although the vectorial model is a reasonable attempt made at simplifying a more accurate but complicated cooperativity such as the one we observe in this work, we did not find any evidence supporting the extent of variation in rates as is suggested by a purely vectorial model.

Our UCG model provides a framework to investigate many important problems related to ATP hydrolysis in actin. Several actin binding proteins (e.g. cofilin (62)) have affinities that depend on the state of the bound nucleotide. In recent work (63), a bottom-up mesoscale modeling approach based on atomistic simulations was used to simulate different modes of applying strain in actin filaments and to study the resulting effect on the binding and activity of the actin binding protein cofilin. Our model can be used to further investigate the implications of applying strain in actin filaments on the hydrolysis of the bound nucleotide and the resulting combined effect of strain and nucleotide state on binding of actin binding proteins.

## AUTHOR CONTRIBUTIONS

HHK, AD and GMH contributed to simulation design and analysis, HHK and AD designed simulation code, HHK, AEPD and AD designed parameter optimization algorithms, GAV designed research. ACS and EMDLC provided crucial biological insights. HHK, AEPD and GMH wrote the manuscript.

## ACKNOWLEDGMENTS

This research was supported in part by the Department of Defense Army Research Office (ARO) through MURI grant W911NF1410403, and also partially supported by the University of Chicago Materials Research Science and Engineering Center (MRSEC), which is funded by National Science Foundation (NSF) under award number DMR-1420709. The computations in this work used the Extreme Science and Engineering Discovery Environment (XSEDE), which is supported by National Science Foundation grant number ACI-1548562. Additional computational resources were provided by the Research Computing Center at The University of Chicago. Glen M. Hocky was supported by a Ruth L. Kirschstein National Research Service Award (NIGMS, F32 GM11345-01). Aleksander E. P. Durumeric acknowledges support by the Department of Defense through the National Defense Science & Engineering Graduate Fellowship (NDSEG) Program.

